# Mitochondria dynamics in postmitotic cells drives neurogenesis through Sirtuin-dependent chromatin remodeling

**DOI:** 10.1101/2020.02.07.938985

**Authors:** Ryohei Iwata, Pierre Vanderhaeghen

## Abstract

The conversion of neural stem cells into neurons is associated with massive remodeling of organelles and chromatin, but whether and how these are linked to control neuronal fate commitment remains unknown. We examined and manipulated mitochondria dynamics with high temporal resolution during mouse and human cortical neurogenesis. We reveal that shortly after cortical stem cells have divided, daughter cells that retain high levels of mitochondria fission will become neurons, while those destined to self-renew undergo rapid mitochondria fusion. Induction of mitochondria fusion after mitosis redirects daughter cells towards cell self-renewal, but only during a restricted time window, which is doubled in human cortical stem cells with higher self-renewing potential. Mitochondria dynamics drives neurogenesis through modulation of the NAD^+^ sensor Sirtuin-1, leading to Histone deacetylation and chromatin remodeling necessary for neurogenic conversion. Our data reveal a post-mitotic critical period of neurogenesis, linking mitochondria state with cell fate.

## Main Text

Neurogenesis is a key event of neural development, by which neural stem/progenitor cells (NSC) stop self-renewing and dividing, to convert to postmitotic neurons. The neurogenic fate transition involves massive changes in shape and polarity of the cell and its organelles (*1*–*3*). Among these, mitochondria dynamics, by which the mitochondria network is remodeled through fusion and fission, has been associated, sometimes causally, with fate changes in various types of stem cells and developing systems, including NSC (*4*–*9*)(*10*)(*11*). On the other hand, as NSC become neurons they undergo global chromatin remodeling, which is required to lead towards stable neuronal fate acquisition, and involves the interplay of neurogenic transcription factors and epigenetic regulators (*12*–*14*).

Organelle changes such as mitochondria fusion/fission are *per se* highly dynamic processes, depending on e.g. cell cycle and metabolic state (*9, 15*), raising the question of how they can be linked to the continuous, irreversible progression towards terminal fate acquisition. Here we examine whether and how the processes of mitochondria and chromatin remodeling are actually coupled, and causally linked, during neurogenesis.

We first looked at mitochondria dynamics and fate acquisition during the neurogenic transition. To this end we first expressed a fluorescent label of mitochondria (mito-GFP) by retroviral infection of radial glia cells (RGC), the major NSC in mouse embryonic cortex, either in vivo or in vitro (Fig. 1A, Fig. S1A). We found that Pax6-positive RGC displayed largely fused mitochondria and Tbr2-positive intermediate progenitors displayed intermediate mitochondrial size, as previously reported (*6*). However, in contrast to previous reports, we also found that early born βIII-tubulin-positive neurons display even shorter, highly fragmented mitochondria, as late as 4 days following their birth (Fig.1A, Fig. S1A), and display mature fused patterns of mitochondria (*16*) only 1-2 weeks afterwards (data not shown). On the other hand, examination of RGC mitochondria during mitosis revealed highly fragmented mitochondria (Fig. S1B), typical of cells in division (*17*), suggesting that mitochondrial dynamics may change in the daughter cells right after mitosis depending on their prospective fate, i.e. as newly born neurons, intermediate progenitors, or RGC.

**Fig. 1.**
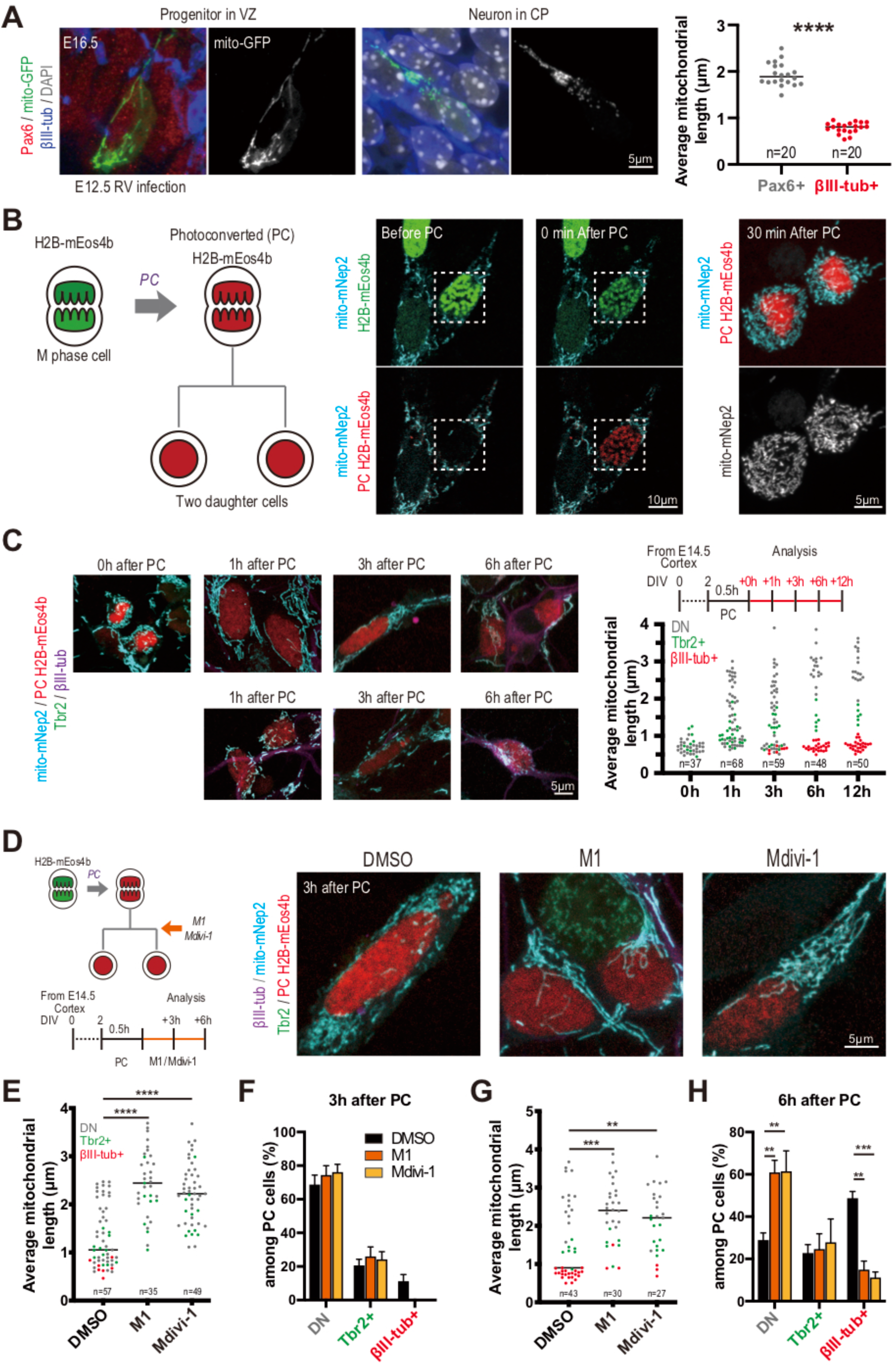
Mitochondria dynamic changes influence fate decision in post-mitotic cortical cells. (A) Representative images of mitochondrial morphology (mito-GFP) in Pax6+ (Apical progenitor in ventricular zone (VZ)) and βIII-tub+ (Newborn neuron in cortical plate (CP)) cell in E16.5 mouse cortex, following in utero retroviral infection at E12.5. Quantified mitochondrial length from two biological replicate experiments. Each data point represents an individual cell average mitochondrial size. ****P < 0.0001; unpaired t test. (B) Schematic of the labeling strategy using photoconverted (PC) histone H2B-mEos4b. Representative images of PC cell labelled with mito-Nep2. (C) Representative images and timeline of PC experiment to determine kinetics of mitochondria dynamics following mitosis in mouse embryonic cortical cells. Quantified mitochondrial length from three biological replicate experiments. Each data point represents an individual cell average mitochondrial size, together with fate marker expression. Red: βIII-tub+ neuron; Green: Tbr2+ intermediate progenitor; Gray: double negative (DN) RGC. (D) Timeline and representative images of PC experiment using M1 and Mdivi-1. (E,G) Quantified mitochondrial length from three biological replicate experiments. (E) 3 hours post-label, (G) 6 hours post-PC. Each data point represents an individual cell average mitochondrial size. **P < 0.01, ***P < 0.001, ****P ≤ 0.0001; Dunnett’s multiple comparisons test. (F,H) Quantification of each cell fate marker+ cells among PC cells from at least four biological replicate experiments. Data are shown as mean ± SEM. **P < 0.01, ***P < 0.001; Dunn’s multiple comparisons test.

To follow mitochondria dynamics of cortical progenitors after cell division with high temporal resolution and cellular throughput, we developed an in vitro assay assessing mitochondrial morphology from cell division to fate acquisition (Fig. 1B). We expressed in cortical progenitors the mitochondrial label mito-mNep2 together with the photoactivatable protein mEOS4b fused to histone H2B. This enabled us to identify unambiguously cells in mitosis, selectively in metaphase/anaphase, based on chromatin mEos labeling, followed by their tagging by photoconversion. The photoconverted (PC) daughter cells could then be tracked from 1 to 24 hours following mitosis, together with their mitochondria. The cells were then fixed and labelled for cell fate markers: βIII-tub for neurons, Tbr2 for intermediate progenitors. The rest of the cells, double-negative (DN) for Tbr2 and βIII-tub, correspond almost exclusively (95%) to Sox2+ RGC (Fig. S1C). Fate acquisition of the daughter cells, towards self-renewing RGC or to neurons, could be observed over the next 6-12 hours (Fig. S1D). Examination of concomitant mitochondrial dynamics revealed distinct patterns in the first 3 hours following cell division, characterized either by mitochondrial length increase over time, or by retaining shorter fragmented mitochondria (Fig. 1C). This pattern correlated remarkably well with later cell fate acquisition, with presumptive RGC displaying long mitochondria, presumptive neurons retaining short mitochondria, and intermediate progenitors displaying intermediate sized mitochondria (Fig. 1C).

As postmitotic alteration of mitochondria dynamics precedes neuronal fate acquisition, could it have a direct influence on neurogenesis? To test this hypothesis, we used the same paradigm of postmitotic cell tracking, but using compounds known to promote mitochondria fusion (M1 (*18*)), or inhibit mitochondria fission (Mdivi-1, Drp1 inhibitor (*19*)). Crucially, the compounds were added to the PC cells right after mitosis, thus enabling to determine the effect of mitochondria dynamics *postmitotically* (Fig. 1D). Treatment with either compound resulted in large increase of mitochondrial size within the first 3 hours of post-mitotic cell labeling (Fig. 1E), as expected, while fate was unaltered at this time-point (Fig. 1F). Examination of the fate of the daughter cells 6 hours after the treatment revealed a dramatic change, with increase in the proportion of RGC, and concomitant decrease of neurons, while the proportion of intermediate progenitors remained unchanged (Fig. 1G,H). This result reflected fate change and not cell loss, as the number of cells was unchanged in either condition after 3-6 hours (Fig. S2A,B).

Examination of the cells 12 hours following M1 treatment revealed a similar increase of RGC at the expense of neurons (Fig. S2C), which was associated with increase of non-neurogenic symmetric divisions at the expense of neurogenic symmetric divisions (Fig. S2D). To examine the self-renewal potential of RGC generated through M1 treatment, we examined cell number and fate of their progeny 24 hours after tracking, thus following further cell division (Fig. S2E). Importantly, this revealed an increase in the number of cell/clones in M1 compared with control conditions, indicating that RGC generated under M1 treatment retained their self-renewal capacity (Fig. S2F,G).

We next determined the upstream mechanisms of post-mitotic mitochondrial dynamics. During mitosis, the mitochondrial fission effector Drp1 is activated by CDK1 phosphorylation (*20*). We examined phospho-Drp1 levels following mitosis during neurogenesis using the mEOS labeling assay. As expected, we found that during mitosis the cells displayed high levels of pDrp1 together with small mitochondria (Fig. S3A). In the following hours however, the cells displayed a dual pattern of phosphorylation, inversely correlated with mitochondrial size changes, with high phosphorylation levels found in the cells with fragmented mitochondria, and low levels found in the cells with large mitochondria (Fig. S3A). We next investigated the effect of an inhibitor of CDK activity, Roscovitine, which as expected could decrease Drp1 phosphorylation and increase mitochondria size in mouse cortical newborn neurons (Fig. S3B). Importantly, postmitotic treatment with Roscovitine also led to concomitant changes in cell fate: we found a decreased proportion of neurons and an increased proportion of RGC, and no detectable cell loss (Fig. S3C-E), further pointing to a crucial role of postmitotic mitochondria dynamics for neuronal fate acquisition.

These data indicate that increased fusion or decreased fission of mitochondria, right after the mother RGC cell division, biases fate acquisition of the daughter cells, which regain stem cell-like fate at the expense of differentiated neuronal fate. As these results were obtained only in vitro, we next examined whether a similar phenomenon could be observed in vivo during mouse corticogenesis. Moreover, as our data so far relied mostly on chemical compounds, to control for the specificity of our findings we first manipulated mitochondria dynamics genetically, using knock-down of Drp1 fission effector through in utero electroporation. This resulted, like for M1 or Mdivi-1 treatment in vitro, in a strong decrease of the generated neurons, while intermediate progenitors and RGC were conversely increased (Fig. 2A,B), confirming the direct implication of mitochondria fission for proper cortical neurogenesis in vivo.

**Fig. 2.**
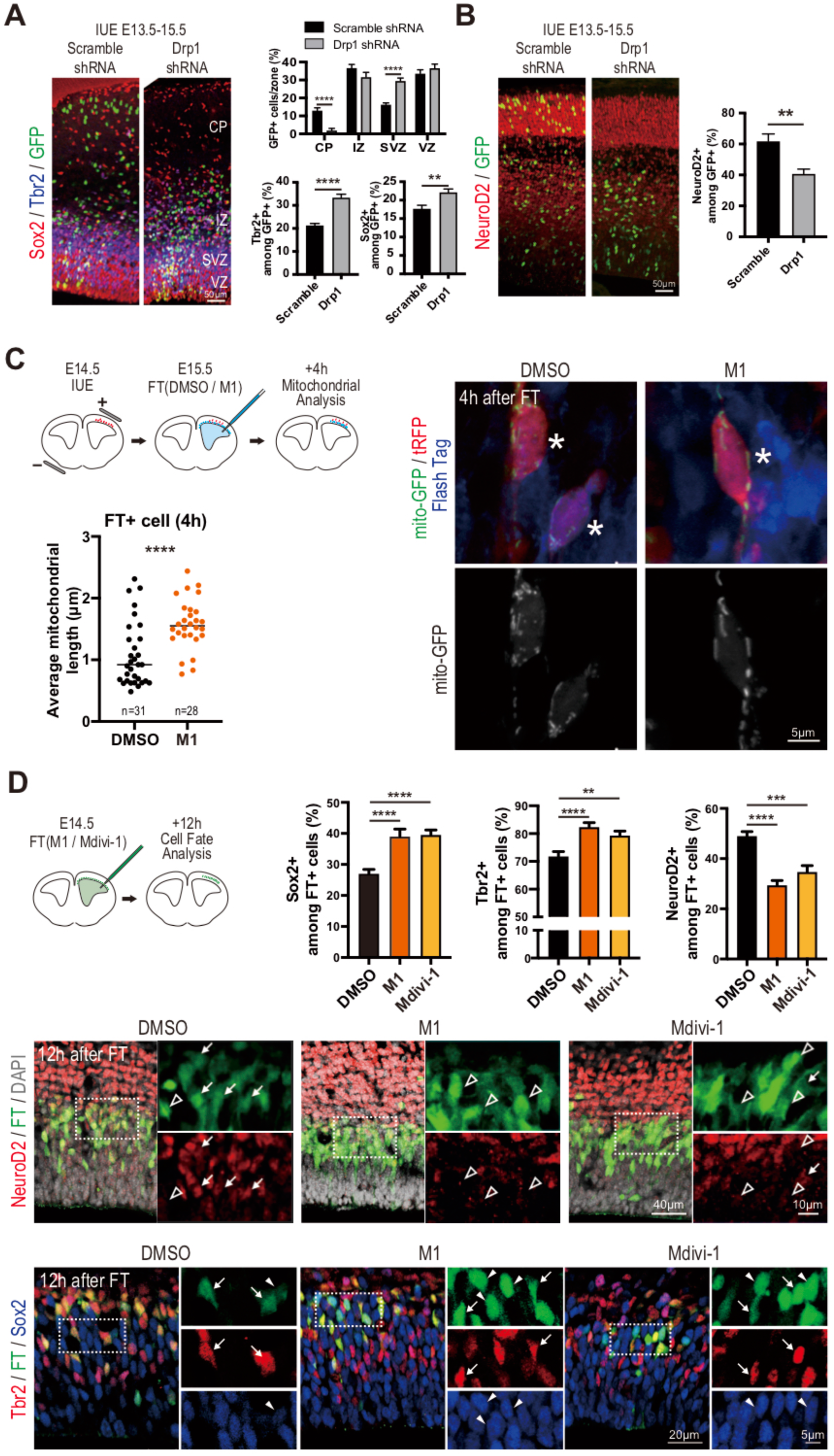
Mitochondria dynamics in post-mitotic cells impact cortical neurogenesis in vivo. (A,B) In utero electroporation (IUE) of scramble or Drp1 shRNA at E13.5, analyzed at E15.5. Histogram show the percentage of H2B-GFP+ cells in VZ, SVZ, IZ, and CP. Quantification of (A) Tbr2+ or Sox2+, and (B) NeuroD2+ cells among electroporated cells from two biological replicate experiments. Data are shown as mean ± SEM. **P < 0.01, ****P < 0.0001; (A, top) Bonferroni’s multiple comparisons test, (A, bottom) Unpaired t test, (B) Mann Whitney test. (C) Timeline of IUE (tRFP and mito-GFP) and Flash Tag (FT) labeling. Representative images of chemical compound treated tRFP and mito-GFP electroporated FT+ cell (Asterisks). Quantification of mitochondrial length from two biological replicate experiments. Each data point represents an individual cell average mitochondrial size. ****P < 0.0001; Mann-Whitney test. (D) Schematic representation and representative images of in utero chemical compound treatment and FT labeling in the mouse embryonic cortex. Quantification of Sox2+, Tbr2+ or NeuroD2+ cells among FT+ cells from two biological replicate experiments. Data are shown as mean ± SEM. **P < 0.01, ***P < 0.001, ****P < 0.0001; (Sox2 and Tbr2) Dunn’s multiple comparisons test, (NeuroD2) Dunn’s multiple comparisons test. Arrow (Top): NeuroD6+, White open arrow head: NeuroD6–, Arrow (Bottom): Sox2+, Arrow head: Tbr2+.

Next we tested specifically whether post-mitotic manipulation of mitochondria dynamics could lead to neural fate changes in vivo. To this aim we took advantage of the FlashTag (FT) fluorescent labeling method (*21*), which enables to label in utero RGC during mitosis, and follow their progeny in vivo. RGC were labelled by in utero injection of FT, together with M1 to promote mitochondria fusion (Fig. 2C). This resulted in increased mitochondria size of the FT+ cells examined 4 hours after labeling and treatment. Remarkably, examination of the FT+ labelled cells 12 hours after M1 or Mdivi-1 treatment revealed an increase in the proportion of Sox2+ RGC and a concomitant decrease of Neurod2+ generated neurons, while the proportion of Tbr2+ intermediate progenitors was slightly increased (Fig. 2D), which could suggest additional inhibitory effects of M1 on conversion of intermediate progenitors into neurons. Overall these data indicate that mitochondria dynamics right after mitosis have a major impact on mouse cortical neurogenesis in vivo, favouring RGC self-renewal and reducing differentiation into neurons, as observed in vitro.

How could mitochondria dynamics impact fate changes early after RGC cell division? One potential mechanism could be through functional changes related to mitochondrial functions (*22*). We first tested the proton ionophore CCCP, which alters the proton gradient necessary for ATP production and thereby leads to overactivation of the oxidative complexes (*23*). This led to a striking increase in neurogenesis within 6 hours post-mitosis (Fig. S4B,C), while the size of the mitochondria appeared to be unchanged over that short period (Fig. S4A).

These data indicate that changes in mitochondrial oxidative activity can promote neuronal fate after RGC mitosis, but given the highly unphysiological mode of action of CCCP, we sought after more biologically relevant targets or effectors. One major sensor of REDOX balance in the cell is the Sirtuin family of deacetylases, through their strong dependence on NAD+/NADH ratio (*24*), which is known to increase following CCCP treatment (*25*). Interestingly Sirtuin-1 (Sirt1) was previously shown to be required for cortical neurogenesis (*26, 27*), where it promotes the suppression of stemness of RGC, through interaction with the BCL6 transcription factor (*26*). BCL6/Sirt1 complex activity leads to chromatin remodeling and thereby to direct transcriptional repression of most key pro-proliferative/stemness genes to promote irreversible fate conversion into neurons (*26, 28*). We therefore tested a potential link between mitochondria function and Sirt1 by testing the impact of post-mitotic Sirt1 blockade on neurogenesis. We first found that Sirt1 inhibition by Ex-527 treatment in post-mitotic cells could effectively block neurogenesis (Fig. 3A, Fig. S4D). Then we tested a potential functional interaction between Sirtuin activity and mitochondria, by assessing the requirement of Sirtuin activity following CCCP treatment. Remarkably, this revealed that CCCP effect on neurogenesis could be completely blocked by Sirtuin inhibition (Fig. S4E,F).

**Fig. 3.**
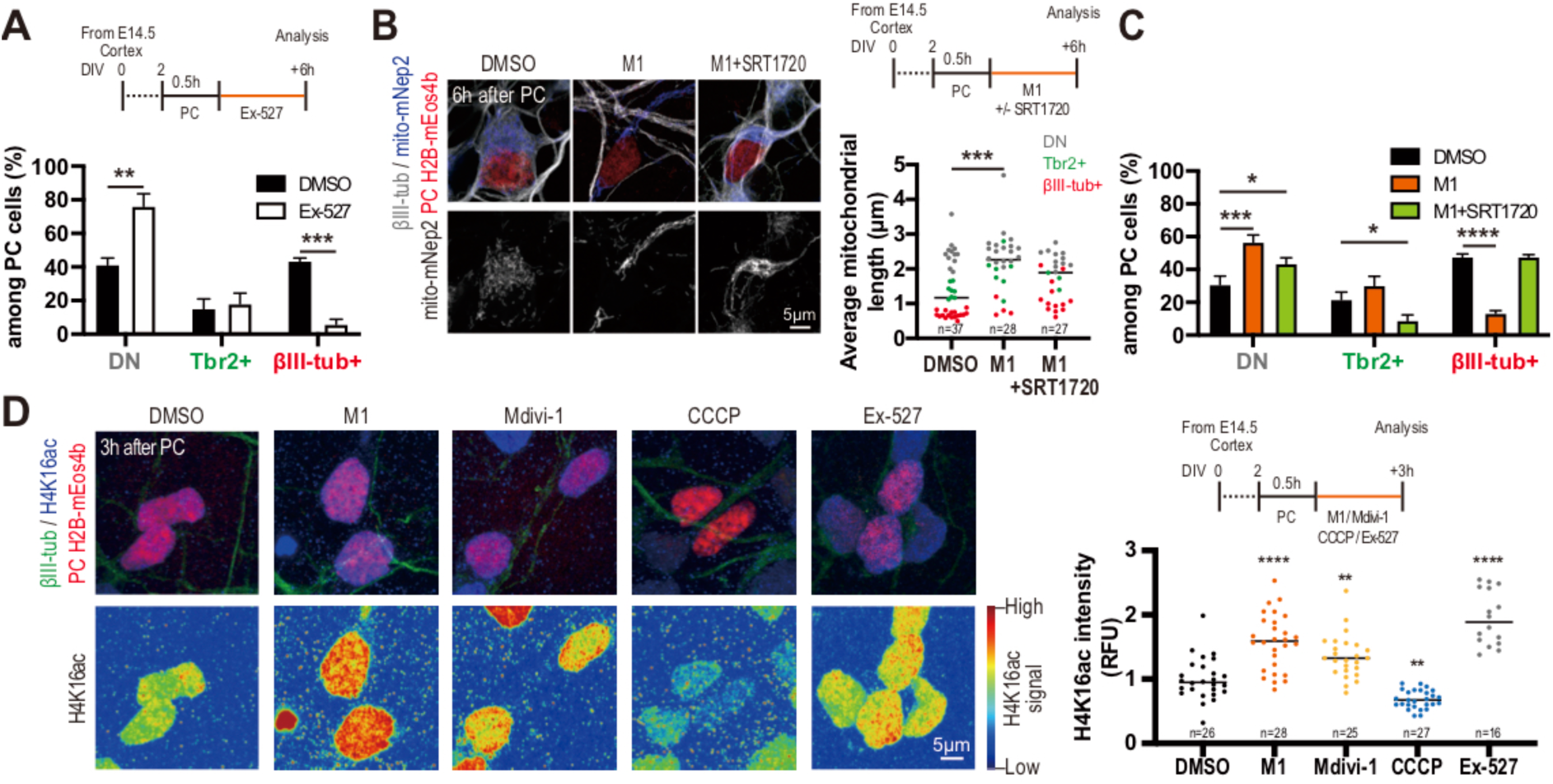
Post-mitotic cell fate determination by mitochondria is mediated through Sirtuin-dependent Histone deacetylation. (A) Timeline and quantification of cell fate marker-positive cells among PC cells using Sirt1 inhibitor Ex-527 from three biological replicate experiments. Data are shown as mean ± SEM. **P < 0.01, ***P < 0.001; Bonferroni’s multiple comparisons test. (B) Timeline and representative images of PC cell using M1 and Sirt1 activator SRT1720. Quantified mitochondrial length from three biological replicate experiments. Each data point represents an individual cell average mitochondrial size. ***P < 0.001; Dunn’s multiple comparisons test. (C) Quantification of each cell fate marker+ cells among PC cells from four biological replicate experiments. Data are shown as mean ± SEM. *P < 0.05, ***P < 0.001, ****P < 0.0001; Dunnett’s multiple comparisons test. (D) Timeline and representative images of H4K16ac signal in PC cells. Quantified H4K16ac signal from two biological replicate experiments. Each data point represents an individual cell average H4K16ac signal. **P < 0.01, ****P < 0.0001; Dunn’s multiple comparisons test.

These data indicate that mitochondria function after mitosis can impact on neurogenesis and that this involves Sirtuin-dependent activity. But does this relate to the mitochondria dynamics, and if so how? To address this crucial point, we tested for a functional interaction between mitochondria fusion and Sirtuin activity, by manipulating mitochondria fusion rates and Sirtuin activity at the same time, right after RGC mitosis. Remarkably, this revealed that the mitochondria-dependent (M1) effects on neurogenesis could be completely abolished by direct Sirtuin activation under SRT1720 treatment (Fig. 3C, Fig. S4G), while Sirtuin activation per se had no effect on mitochondria morphology (Fig. 3B).

These data thus indicate that M1-mediated mitochondria fusion inhibits neuronal differentiation through modulation of NAD^+^-dependent Sirtuin activity. On the other hand, it was previously shown that Sirtuins act during neurogenesis through major changes in chromatin remodelling, in particular through H4K16 deacetylation of BCL6 transcriptional targets, thereby leading to their repression (*26, 28*). We therefore examined the global pattern of H4K16 acetylation in response to M1, Mdivi-1 or Ex-527 in the hours following mitosis of RGC. This revealed a striking global increase of H4K16 acetylation levels, further indicating that these treatments have a direct impact on chromatin remodeling during neurogenesis. On the other hand, H4K16 acetylation levels were decreased through CCCP treatment, further linking nuclear Sirtuin activity to mitochondria OXPHOS activity (Fig. 3D). Overall these data indicate that mitochondrial dynamics exert its influence on early post-mitotic fate determination at least in a large part through Sirtuin modulation, which thereby leads to chromatin remodeling instrumental for neurogenic fate commitment.

Human cortical progenitors are characterized by increased self-renewal potential that enables their expansion and ultimately increased neuronal output, which is thought to underlie the increase in cortical size in the human lineage (*29, 30*). We therefore examined patterns of mitochondria dynamics during in vitro corticogenesis from human pluripotent stem cells, a system in which such human-specific features are recapitulated (*31, 32*). As in the mouse, human cortical RGC in interphase were characterized by large sized mitochondria, while early born neurons displayed fragmented mitochondria (Fig. 4A). Moreover and importantly, the overexpression of mitochondria fission-promoting Drp1 or MFF genes in human RGC resulted in a strong increase in neurogenesis (Fig, 4B), further suggesting that the mechanisms uncovered in the mouse were conserved in human corticogenesis.

**Fig. 4.**
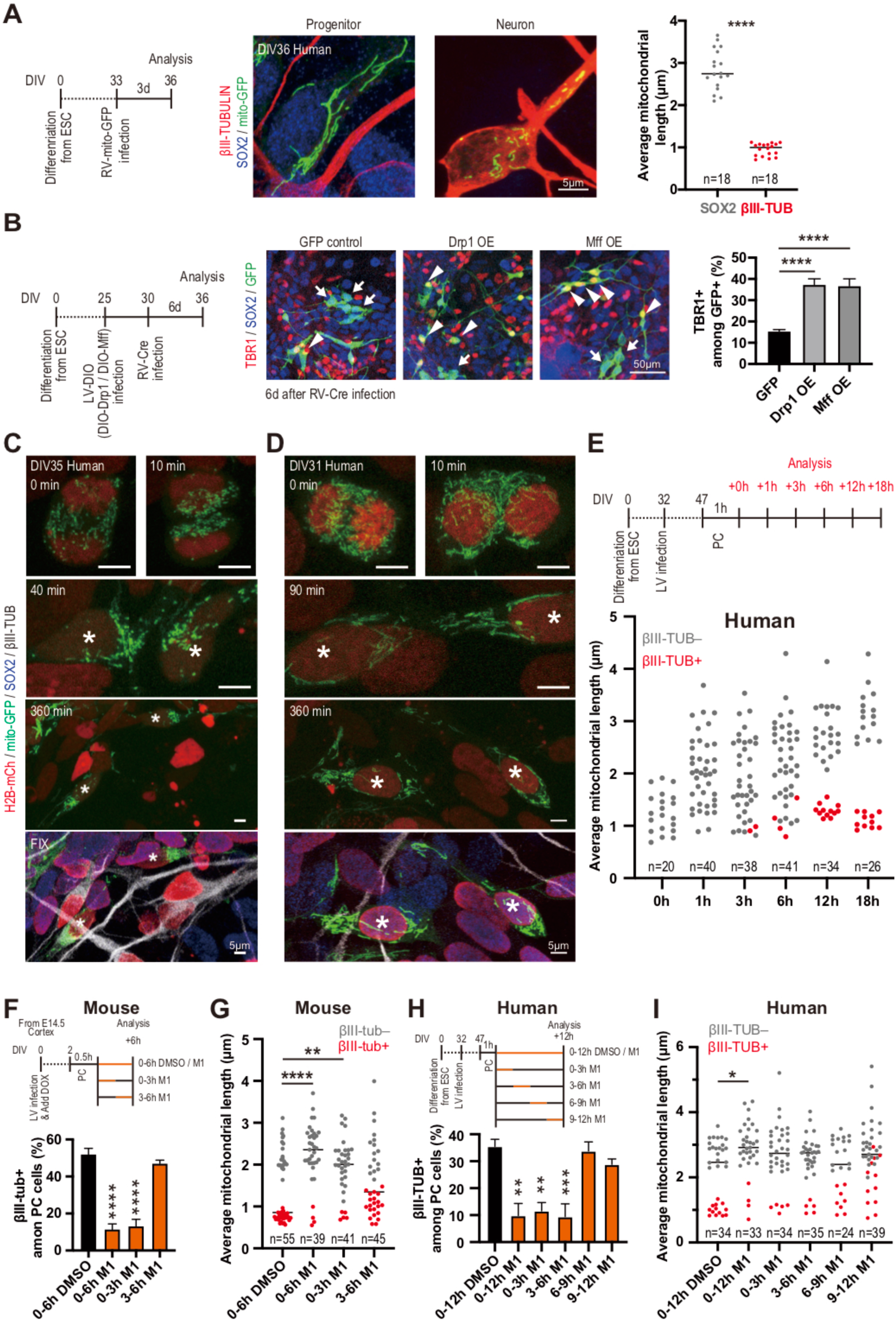
A species-specific post-mitotic critical period for neuronal fate commitment. (A) Representative images of mitochondrial morphology in SOX2+ (RGC) and βIII-tub+ (Newborn neuron) human ESC-derived cortical cell 3 days post mito-GFP retroviral infection. Quantification of mitochondrial length from two biological replicate experiments. Each data point represents an individual cell average mitochondrial size. ****P < 0.0001; Mann Whitney test. (B) Timeline and representative images of Drp1- and Mff-overexpressing human cortical cells (6 days post Cre-expressing retrovirus infection). Arrow head: Neuron (Tbr1+), Arrow: Progenitor (Sox2+). Quantification of TBR1+ cells among GFP-labeled cells from three biological replicate experiments. Data are shown as mean ± SEM. ****P < 0.001; Dunnett’s multiple comparisons test. (C,D) Representative time-lapse images of mitochondrial dynamics in (C) Symmetric non-neurogenic division and (D) Symmetric neurogenic division. Asterisks indicate tracked cells. (E) Timeline of PC experiment and quantified mitochondrial length from three biological replicate experiments. Each data point represents an individual cell average mitochondrial size. (F,H) Timing of M1 treatment in (F) mouse and (H) human cortical cells. Quantification of βIII-tub+ cells among PC cells from three biological replicate experiments. Data are shown as mean ± SEM. **P < 0.01, ***P < 0.001, ****P < 0.0001; Dunnett’s multiple comparisons test. (G,I) Quantification of mitochondrial length in mouse (G) and human (I) cortical cells from three biological replicate experiments. Each data point represents an individual cell average mitochondrial size. *P < 0.05, **P < 0.01, ****P < 0.0001; Dunn’s multiple comparisons test.

To test this further we performed time-lapse imaging of human cortical progenitors labelled with mito-GFP, followed by post hoc fate marker determination (Fig. 4C,D). This revealed that at mitosis the mitochondria were fragmented, followed by a dual pattern similar to the mouse, with large mitochondria sized cells remaining progenitors, while those destined to become neurons kept fragmented mitochondria (n=24 cells). Similar data were obtained using mEOS labeling of human mitotic RGC, with increased mitochondria size found in the self-renewing progenitors and decreased mitochondria size among newly born neurons, like in the mouse (Fig. 4E, fig. S5A). Moreover, M1 treatment following mitosis of human progenitors was found to lead to increased mitochondria size and decreased neuronal differentiation, as well as increased self-renewing division types (Fig. 4I, Fig. S5B,C).

These data indicate that the postmitotic control of fate conversion through mitochondria dynamics is overall conserved between mouse and human corticogenesis. But given the differences in the balance between self-renewal and neurogenesis in mouse vs. human progenitors, we then determined using the mEOS system the length of the “critical” period during which mitochondria dynamics can actually impact neural cell fate. Indeed, given the higher self-renewal potential of human RGC, one could speculate that the critical period of plasticity enabling to favour self-renewal, is longer in these cells. Human and mouse cells were thus treated in parallel over restricted time-periods following mitosis (Fig. 4F-I). Remarkably, this revealed that while in the mouse M1 treatment could alter cell fate until 3 hours post-mitosis, but not beyond (Fig. 4F), its impact on human cells could be detected for at least 6 hours after mitosis (Fig. 4H). These data indicate that the critical period of post-mitotic neural cell fate plasticity appears to be doubled in the human progenitors, in line with their increased self-renewal potential.

The mechanisms that determine the timing of cell fate commitment remain a central unanswered question in stem cell and developmental biology. Previous data emphasized on fate plasticity and commitment of neural stem cell before mitosis, as early as G1 in the mother cell (*33*–*36*), with post-mitotic neuronal cells becoming increasingly resilient to extrinsic or intrinsic cues (*36*). Our data rather indicate the existence of a key fate decision point much later, i.e. in daughter cells shortly after NSC mitosis. This is reminiscent of embryonic stem cell differentiation that is also thought to occur in G1 (*37, 38*), especially early G1 at a stage most favorable to massive chromatin remodeling (*39*). Moreover, we find that this critical period is doubled in human cortical progenitors, which could contribute to their increased self-renewal capacities, thus uncovering a surprising link between mitochondria dynamics and human brain evolution.

The conversion of NSC into intermediate neural progenitors is associated with increased mitochondria fission and oxidative activity (*6, 40*), and neuronal differentiation itself is accompanied by increased oxidative metabolism (*41*–*45*), but the causal relationships between these events, and neuronal fate acquisition itself, had remained unclear. Our data rather indicate that during mouse and human cortical neurogenesis, newly born neurons *themselves* display surprisingly high levels of fission, which then lead to their irreversible fate commitment, through REDOX-sensitive Sirtuin-dependent chromatin remodeling.

How mitochondria shape and function are linked during neurogenesis remains to be explored further, but our data indicate that mitochondria dynamics biases neural cell fate choice through increased NAD+-dependent Sirtuin activity, specifically in early post-mitotic phase. Interestingly this phase is also characterized by lowest levels of expression of Hes Notch effectors (*46*), which is directly repressed by Sirt1 (*26, 27*). As Sirtuin activity has been shown to modulate stem cells in other systems (*47*), this raises the possibility that the mitochondria dynamics-dependent modulation of Sirtuin activity shortly after mitosis could contribute linking cell state with fate changes in other stem cells and developing systems, as well as during neuronal direct reprogramming (*48*).

## Acknowledgments

The authors thank members of the Vanderhaeghen lab and CBD for helpful discussions, and J.-M. Vanderwinden of the ULB Light Microscopy Facility for his support with imaging. Some of the images were acquired on a Zeiss LSM 880 – Airyscan system (Cell and Tissue Imaging Cluster, CIC), supported by Hercules AKUL/15/37_GOH1816N and FWO G.0929.15 to Pieter Vanden Berghe, KU Leuven.

## Author contributions

Conceptualization and Methodology, R.I. and P.V.; Investigation, R.I. and P.V.; Formal Analysis, R.I. and P.V.; Writing - R.I. and P.V; Funding acquisition, P.V.; Resources, P.V.; Supervision, P.V. Authors declare no competing interests. All data is available in the main text or the supplementary materials.

## Materials and Methods

### Mice

All mouse experiments were performed with the approval of the KU Leuven and the Université Libre de Bruxelles Committees for animal welfare. Mice were housed under standard conditions (12h light:12h dark cycles) with food and water ad libitum. Embryos (aged E12.5 - E16.5) of the mouse strain ICR (CD1, Charles River Laboratory) were used for in utero experiments and primary culture. The plug date was defined as embryonic day (E)0.5, and the day of birth was defined as P0. The data obtained from all embryos were pooled without discrimination of sexes for the analysis of in utero electroporation, given the difficulty to determine sex identity at embryonic stages.

### DNA Construct

The DNA fragment of H2B-mEos4b-6 (a gift from Michael Davidson (Addgene plasmid # 57508; http://n2t.net/addgene:57508; RRID:Addgene_57508)) and TRE (pTRE-tight vector (Clontech #631059)) were prepared by PCR and transferred to lentiviral plasmid backbone (a gift from Cecile Charrier) by restriction digestion and ligation to obtain a pLenti-TRE-H2B-mEos4b-WPRE. DNA fragment of mito-mNep2-FLAG (mNeptune2-N1, a gift from Michael Davidson (Addgene plasmid # 54837; http://n2t.net/addgene:54837; RRID:Addgene_54837), Mitochondrial target sequence with linker in N-terminal: ATGTCCGTCCTGACGCCGCTGCTGCTGCGGGGCTTGACAGGCTCGGCCCGGCGGCTCCCAGTGCCGCGCGCCAAGATCCATTCGTTGGGGGATCCACCGGTCGCCACC, FLAG in C-terminal: GATTACAAGGATGACGACGATAAG) were prepared by PCR and transferred to CAG promoter (a gift from Hitoshi Niwa) containing lentiviral plasmid backbone by restriction digestion and ligation to obtain a pLenti-CAG-mito-mNep2-FLAG-WPRE.

The DNA fragment of LoxP-Stop-LoxP (a gift from Takuji Iwasato (Addgene plasmid # 69138; http://n2t.net/addgene:69138; RRID:Addgene_69138)) and mito-EmGFP (a gift from Michael Davidson (Addgene plasmid # 54160; http://n2t.net/addgene:54160; RRID:Addgene_54160)) were transferred to TRE promoter lentiviral plasmid backbone by restriction digestion and ligation to obtain a pLenti-TRE-LSL-mito-EmGFP-WPRE. DNA fragment of DIO-EGFP (a gift from Bernardo Sabatini (Addgene plasmid # 37084; http://n2t.net/addgene:37084; RRID:Addgene_37084)) was transferred to CAG promoter containing lentiviral plasmid backbone by restriction digestion and ligation to obtain a pLenti-CAG-DIO-GFP-WPRE. DNA fragment of mDrp1-P2A (a gift from David Chan (Addgene plasmid # 34706; http://n2t.net/addgene:34706; RRID:Addgene_34706), P2A sequence in C-terminal: GCTACTAACTTCAGCCTGCTGAAGCAGGCTGGAGACGTGGAGGAGAACCCTGGACCT) was prepared by PCR and transferred to CAG-DIO lentiviral plasmid backbone by restriction digestion and ligation to obtain a pLenti-CAG-DIO-mDrp1-P2A-GFP-WPRE. DNA fragment of mMff-P2A (a gift from David Chan (Addgene plasmid # 44601; http://n2t.net/addgene:44601; RRID:Addgene_44601), P2A sequence in C-terminal: GCTACTAACTTCAGCCTGCTGAAGCAGGCTGGAGACGTGGAGGAGAACCCTGGACCT) was prepared by PCR and transferred to CAG-DIO lentiviral plasmid backbone by restriction digestion and ligation to obtain a pLenti-CAG-DIO-mMff-P2A-GFP-WPRE. DNA fragment of mito-EmGFP and PGK promoter (a gift from Tyler Jacks (Addgene plasmid # 11586; http://n2t.net/addgene:11586; RRID:Addgene_11586)) were prepared by PCR and transferred to a retroviral plasmid backbone (a gift from Fred Gage (Addgene plasmid # 49054; http://n2t.net/addgene:49054; RRID:Addgene_49054)) by restriction digestion and ligation to obtain a pRetro-PGK-mito-EmGFP-WPRE. DNA fragment of Cre was prepared by PCR and transferred to retroviral plasmid backbone (a gift from Fred Gage (Addgene plasmid # 49054; http://n2t.net/addgene:49054; RRID:Addgene_49054)) by restriction digestion and ligation to obtain a pRetro-CAG-Cre-WPRE. DNA fragment of H2B-EmGFP was prepared by PCR and transferred to the pSCV2 backbone (a gift from Franck Polleux) with the Drp1 shRNA sequence (CGTATCAGTCTCTTCTAAATGCTTCCTGTCACATTTAGAAGAGACTGATACTTTTTT) by restriction digestion and ligation to obtain a pCAG-H2B-EmGFP-U6-Drp1 shRNA.

### Lentiviral preparation

HEK293T cells were transfected by packaging plasmids, psPAX2 (a gift from Didier Trono (Addgene plasmid # 12260; http://n2t.net/addgene:12260; RRID:Addgene_12260)) and pMD2.G (a gift from Didier Trono (Addgene plasmid # 12259; http://n2t.net/addgene:12259; RRID:Addgene_12259)), and a plasmid of gene of interest in lentiviral backbone (pLenti-TRE-H2B-mEos4b-WPRE, pLenti-CAG-mito-mNep2-FLAG-WPRE, pLenti-DIO-CAG-EGFP-WPRE, pLenti-DIO-CAG-mDpr1-P2A-EGFP-WPRE, pLenti-DIO-mMff-P2A-EGFP-WPRE, pLenti-PGK-H2B-mCherry-WPRE (a gift from Mark Mercola (Addgene plasmid # 21217; http://n2t.net/addgene:21217; RRID:Addgene_21217)), and FUW-M2rtTA (a gift from Rudolf Jaenisch (Addgene plasmid # 20342; http://n2t.net/addgene:20342; RRID:Addgene_20342)). Three days after transfection, culture medium was collected and viral particles were enriched by filter device (Amicon Ultra-15 Centrifuge Filters, Merck, Cat#UFC910008). Titer check was performed on HEK293T cell culture for every batch of lentiviral preparation.

### Retrovirus preparation

HEK293T cells were transfected by packaging plasmids, pCMV-VSV-G (Addgene plasmid # 8454) and pUMVC (Addgene plasmid # 8449), and a plasmid of gene of interest in a retroviral backbone (pRetro-PGK-mito-EmGFP-WPRE, and pRetro-CAG-Cre-WPRE).

Three days after transfection, culture medium was collected and viral particles were enriched by filter device (Amicon Ultra-15 Centrifuge Filters, Merck, Cat#UFC910008). Titer check was performed on HEK293T cell culture for every batch of lentiviral preparation.

### Primary mouse cortical cell culture

Timed pregnant mouse embryos at E13.5-E14.5 were used. Cerebral cortices were dissected and enzymatically dissociated using Trypsin-EDTA (Thermo Fisher) with DNase I (VWR). The tissue was triturated with a glass Pasteur pipette to generate a single-cell suspension. The single cells were plated on laminin- and poly-D-lysine-coated coverslips or coverslip bottom dishes with grid (ibidi, Cat#81166) at high confluency (300,000 cells/cm2) using culture medium: Neurobasal (Thermo Fisher, Cat#21103049) with GlutaMAX supplement (1x, Thermo Fisher, Cat# 35050061), sodium pyruvate (1mM, Thermo Fisher, Cat#11360070), Penicillin-Streptomycin (50 U/ml, Thermo Fisher, Cat#15070063), B-27 supplement (1x, Thermo Fisher, Cat# 17504044), N-2 supplement (1x, Thermo Fisher, Cat#17502048), N-acetyl-cysteine (1mM, Sigma-Aldrich, Cat#A7250), bFGF (10 ng/ml, PeproTech) and Doxycycline hyclate (1 mg/ml, Sigma-Aldrich, Cat#D9891). A mixture of lentiviral vectors was added to cortical cells for infection. The next day, medium was changed to Neurobasal with GlutaMAX supplement, Penicillin-Streptomycin, B-27 supplement and Doxycycline. Plated cells were incubated at 37°C with 5% CO2.

### Human ESC culture and cortical cell differentiation

Human ESC (H9; WiCell Cat # NIHhESC-10-0062; female donor) was grown on irradiated mouse embryonic fibroblasts in the ES cell medium until the start of cortical cell differentiation. Cortical cell differentiation was performed as described previously (*1, 2*) and frozen at day 25. Differentiated cortical cells were validated for neuronal and cortical markers by immunostaining using antibodies for TUBB3 (BioLegend, Cat#MMS-435P), TBR1 (Abcam, Cat#ab183032), CTIP2 (Abcam, Cat#ab18465), FOXG1 (Takara, Cat#M227), Sox2 (Santa Cruz, Cat#sc-17320), FOXP2 (Abcam, Cat#ab16046), SATB2 (Abcam, Cat#ab34735), and CUX1 (Santa Cruz, Cat#sc-13024).

### Human cortical cell culture

Human cortical cells (frozen at day 25) were thawed and plated on Matrigel-coated (BD, Cat#354277) plates using DDM/B27 and Neurobasal supplemented with B27 (DDM/B27+Nb/B27) medium (*1, 2*) at 37°C with 5% CO2. For the mitochondrial labeling experiment (Fig. 4A), at five days after thawing, cells were dissociated using Accutase (Thermo Fisher, Cat#00-4555-56) and plated on mouse astrocyte coated coverslips at low confluency (20,000-100,000 cells/cm2). At three days after co-culture with astrocytes, cells were infected with mito-EmGFP-expressing retroviral vector. The next day, medium was changed to DDM/B27+Nb/B27 medium. At three days after retroviral vector infection, coverslips were fixed with 4% paraformaldehyde in PBS. For the time-lapse experiments (Fig. 4C,D), H2B-mCherry-expressing lentiviral vector was infected at day 25. At three days after lentivirus infection, cells were dissociated using Accutase plated on mouse astrocyte coated coverslip bottom dish at low confluency (20,000-100,000 cells/cm2) with mito-EmGFP-expressing retroviral vector. The next day, medium was changed to DDM/B27+Nb/B27 medium. For the cell-tracking experiments with mitochondrial morphology labeling (Fig. 4E,H,I, fig. S5A), seven days after thawing, cells were dissociated using Accutase and plate on mouse astrocyte coated coverslips at high confluency (500,000-900,000 cells/cm2) with H2B-mEos4b-expressing and rtTA-expressing lentiviral vector. The next day, medium was changed to DDM/B27+Nb/B27 medium. Ten days after lentiviral vector infection, cells were dissociated using Accutase and plated on mouse astrocyte coated coverslips bottom dish at high confluency (300,000 cells/cm2). At three days after coculture with astrocyte, medium was changed to phenol red minus DDM/B27+Nb/B27 medium with Doxycycline (1 mg/ml, Sigma-Aldrich). For the 12h M1 treatment experiment (fig. S5B,C), seven days after thawing, cells were dissociated using Accutase and plated on mouse astrocyte coated coverslips at high confluency (500,000-900,000 cells/cm2) with H2B-mEos4b-expressing and rtTA-expressing lentiviral vector. The next day, medium was changed to DDM/B27+Nb/B27 medium. Three days after lentiviral vector infection, cells were dissociated using Accutase and plated on mouse astrocyte coated coverslips bottom dish at high confluency (300,000 cells/cm2). Three days after coculture with astrocyte, medium was changed to DDM/B27+Nb/B27 medium with Doxycycline. For the overexpression experiment (Fig. 4B), DIO-lentiviral vector (DIO-GFP, DIO-Drp1-P2A-GFP, or DIO-Mff-P2A-GFP) was infected at day 25. At three days after lentivirus infection, cells were dissociated using Accutase and plated on mouse astrocyte coated coverslip at high confluency (100,000 cells/cm2). Cre-expressing retroviral vector was infected two days after coculture with astrocytes. Six days after retroviral vector infection, coverslips were fixed with 4% paraformaldehyde in PBS.

### Imaging and photoconversion

The medium was changed to phenol red-free Nb/B27 (Mouse) or DDM/B27+Nb/B27 (Human) medium before imaging. During imaging, cells were maintained at 37°C under humidified air with 5% CO2. The mitotic cells (prometaphase, metaphase, and anaphase) were identified using histone H2B-mEos4b signal and photoconverted by laser stimulation at 405 nm. After photoconversion, medium was changed to Nb/B27 (Mouse) or DDM/B27+Nb/B27 (Human) medium with following chemical compounds: M1 (50µM, Sigma-Aldrich, Cat#SML0629), Mdivi-1 (50µM, Sigma-Aldrich, Cat#M0199), Roscovitine (50µM, Sigma-Aldrich, Cat#R7772), CCCP (10µM, Abcam, Cat#ab141229), Ex-527 (50µM, Santa Cruz Biotechnology, Cat#sc-203044), or SRT1720 (2µM, Santa Cruz Biotechnology, Cat#sc-364624).

### Analysis of mitochondrial morphology

Morphologies were imaged on a Zeiss LSM780 or LSM880 confocal microscope, using 40x (NA 1.3) or 63x (NA 1.4) oil immersion objectives with 4.0x digital zoom, at a horizontal resolution of 0.088–0.138 µm/pixel and a vertical resolution of 0.2–0.4 µm/pixel. For each cell, a z-stack of 10-20 images was acquired centered on the cell of interest. A maximal intensity projection image was performed. An image of the first cell was thresholded visually as to optimally individualize mitochondria fragments using raw mitochondrial membrane signal (e.g. mito-GFP, mito-mNep2). This image was later used as a “reference” image for visual thresholding of the other cell images. Mitochondria fragments were defined as belonging to a cell based on the distance from the nucleus (e.g. H2B-mEos4b signal) and the difference of mitochondrial membrane signal intensity between neighbouring cells. The freehand tool in Fiji was used to manually trace the maximum length of mitochondria fragment. The 3D projection tool or serial images were used to determine whether there is a connection between mitochondria fragments. Only mitochondria fragments which could be visually identified as single mitochondria were measured.

### Analysis of relative fluorescence units

To quantify global p-Drp1 and H4K16ac protein levels in PC cortical cells (fig. S3A and Fig. 4D), fluorescence images were acquired on a Zeiss LSM780 confocal microscope, using 63x (NA 1.4) oil immersion objectives, at a horizontal resolution of 0.088–0.138 µm/pixel and a vertical resolution of 0.2–0.4 µm/pixel under identical conditions. The maximum intensity projection images were used for measuring signal intensity by Fiji software. The final value of relative fluorescence units (RFU) was determined for each condition using the following calculation:

(Region of interest of signal intensity) – (Area of background staining of signal intensity)

For calculating the RFU of H4K16ac (Fig. 4D), each signal intensity was normalized by mean value of signal intensity of DMSO treated cell condition.

### In utero retrovirus injection

For in vivo mitochondrial labeling (Fig. 1A), timed-pregnant mice were anesthetized with a mixture of ketamine (Ketalar, 50mg/ml solution injectable, Pfizer) and xylazine (Rompun, 2% solution injectable, Bayer) at E12.5. The mito-EmGFP-expressing retroviral solution was injected into the lateral ventricles of the embryos using a heat-pulled capillary. Embryos were placed back into the abdominal cavity, and mice were sutured and placed on a heating plate until recovery.

### In utero electroporation

In utero electroporation was performed as described previously (*3*). Timed-pregnant mice were anesthetized with a mixture of ketamine (Ketalar, 50mg/ml solution injectable, Pfizer) and xylazine (Rompun, 2% solution injectable, Bayer) or Isoflurane (Iso-Vet, 1000 mg/g, Dechra) at E13.5. Plasmid solutions containing 0.1-1.0 µg/µl each of DNA were injected into the lateral ventricles of the embryos using a heat-pulled capillary. Electroporation was performed using tweezer electrodes (Nepa Gene, Cat#CUY650P5) connected to a BTX830 electroporator (five pulses of 25-40 V for 100 ms with an interval of 1 s). Embryos were placed back into the abdominal cavity, and mice were sutured and placed on a heating plate until recovery. For the mitochondrial labeling experiment (Fig. 2C), DNA mixture of pCAG-LSL-tRFP-ires-tTA-WPRE (1.0 µg/µl), pCAG-Cre-WPRE (100 ng/µl), and pTRE-LSL-mito-EmGFP-WPRE (1.0 µg/µl) were electroporated at E14.5. Twenty-four hours later, 0.5 µl of Far Red CellTraceTM (0.5 mM in DMSO, Thermo Fisher, Cat#C34564) with or without M1 (25mM, Sigma-Aldrich, Cat#SML0629) were injected in the electroporated side ventricle. These embryos were sacrificed four hours after Flash Tag labeling. For the knockdown experiment (Fig. 2A,B), DNA of pCAG-H2B-EmGFP-U6-Drp1 shRNA (1.0 µg/µl) was electroporated at E13.5.

### Flash Tag

Flash Tag labeling was performed as described previously (*4*) with some modification. Timed-pregnant mice were anesthetized with Isoflurane (Iso-Vet, 1000 mg/g, Dechra) at E14.5. CFSE CellTraceTM (2.5 mM in DMSO, Thermo Fisher, Cat#C34554) together with M1 (25mM, Sigma-Aldrich, Cat#SML0629) or Mdivi-1 (50mM, Sigma-Aldrich, Cat#M0199) were injected into the lateral ventricles of the embryos using a heat-pulled capillary. These embryos were sacrificed twelve hours after Flash Tag labeling.

### Immunostaining

For immunocytochemistry, coverslips were fixed with 4% PFA in PBS 1h at 4°C. The coverslips were transferred into PBS, then blocked PBS with 3% horse serum (Thermo Fisher) and 0.3% Triton X-100 (Sigma-Aldrich, Cat#T9284) during 1 h, and incubated overnight at 4°C with the primary antibodies. After three washes with PBS/0.1% Triton X-100, slices were incubated in PBS for 5 min at room temperature and incubated 2 h at room temperature with the appropriated secondary antibodies with DAPI (Sigma-Aldrich, Cat#D9542). Sections were again washed three times with PBS for 5 min. The sections were mounted on a Superfrost slide (Thermo Fisher) and add Glycergel mounting medium (Dako, Cat#C0563). For immunofluorescence section, mice were fixed with 4% PFA in PBS. Brains were dissected, and 100 µm sections were prepared using a Leica VT1000S vibratome. Slices were transferred into PBS with sodium azide (0.5 µg/mL: Sigma-Aldrich, Cat#71289), then blocked with PBS with 3% horse serum and 0.3% Triton X-100 during 1 h, and incubated two overnight at 4°C with the primary antibodies. After three washes with PBS/0.1% Triton X-100, slices were incubated in PBS for 1 h at room temperature and incubated 2 h at room temperature with the appropriated secondary antibodies with DAPI. Sections were again washed three times with PBS for 1 h. The sections were mounted on a Superfrost slide (Thermo Fisher) and add Glycergel mounting medium. The cells were immunostained using antibodies for Mitochondria (Abcam, Cat#ab92824, GFP (Abcam, Cat#ab13970), mCherry (Thermo Fisher, Cat#M11217), DDDDK (Abcam, Cat#ab1257), Phospho-Drp1(Ser616) (Cell signaling, Cat#3455S), Phospho-Histone H3 (Ser28) (Abcam, Cat#ab10543), TUBB3 (BioLegend, Cat#MMS-435P), TUBB3 (BioLegend, Cat#PRB-435P), Tbr1 (Abcam, Cat#ab183032), Pax6 (BD, Cat#561462), Sox2 (Santa Cruz, Cat#sc-17320), NeuroD2 (Abcam, Cat#ab104430), Tbr2 (Abcam, Cat#ab183991), and Histone H4 (acetyl K16) (Abcam, Cat#ab109463). Imaging was performed using either a Zeiss LSM780 or LSM880 confocal microscope controlled by the Zen Black software (Zeiss).

### Statistical analysis

All statistical analyses were performed using GraphPad Prism 8 (GraphPad Softwares). Shapiro-Wilk normality test was used to test the normality. Parametric data were analyzed by t-test, one-way or two-way ANOVA followed by Dunnett’s, or Bonferroni’s post hoc analysis for comparisons of multiple samples. Non-parametric data were analyzed by the Mann-Whitney test or Kruskal-Wallis test followed by Dunn’s post hoc analysis for comparisons of multiple samples. P values < 0.05 were considered statistically significant. Data are presented as mean ± SEM or median.

**Fig. S1.**
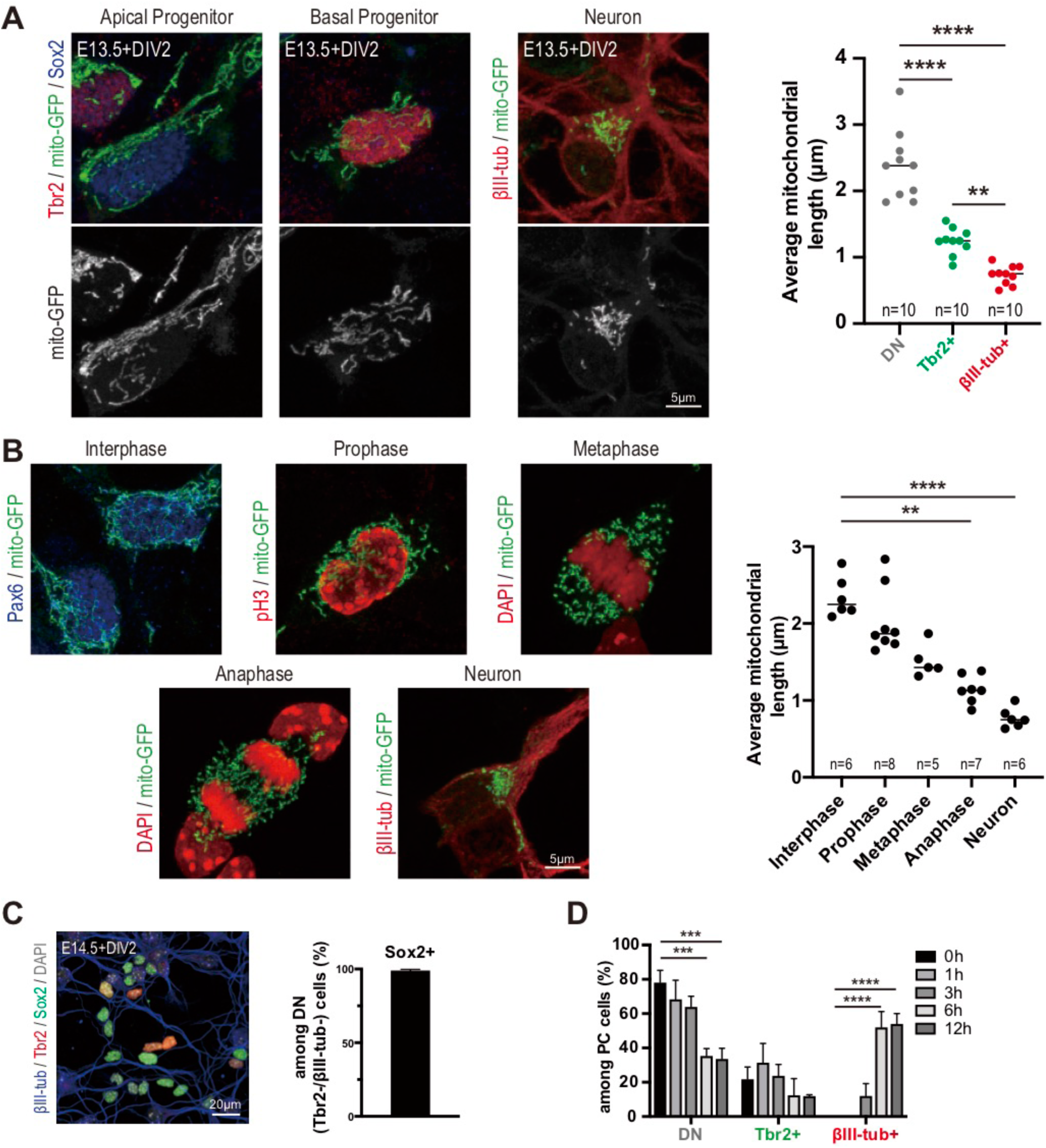
Mitochondria dynamics in mouse corticogenesis in vitro. (A) Representative images of mitochondrial morphology (mito-GFP) in Pax6+ (RGC), Tbr2+ (Intermediate progenitor), and βIII-tub+ (Neuron) cells in vitro (E13.5 cortex cultured for 2 days in vitro (DIV)). Quantified mitochondrial length from two biological replicate experiments. Each data point represents an individual cell average mitochondrial size. **P < 0.01, ****P < 0.0001; Bonferroni’s multiple comparisons test. (B) Representative images of mitochondrial morphology during mitosis of cortical progenitors in vitro (E13.5 cortex cultured for 2 DIV). Quantified mitochondrial length. Each data point represents an individual cell average mitochondrial size. **P < 0.01, ****P < 0.0001; Dunn’s multiple comparisons test. (C) Representative images of Sox2+ (RGC), Tbr2+ (Intermediate progenitor), and βIII-tub+ (Neuron) cells (E14.5 cortex cultured for 2 DIV). Quantification of Sox2+ cells among DN (Tbr2–/βIII-tub–) cells from two biological replicate experiments. (D) Quantification of each cell fate marker+ cells among PC cells from three biological replicate experiments (cf. Fig.1C). Data are shown as mean ± SEM. ***P < 0.001, ****P < 0.0001; Bonferroni’s multiple comparisons test.

**Fig. S2.**
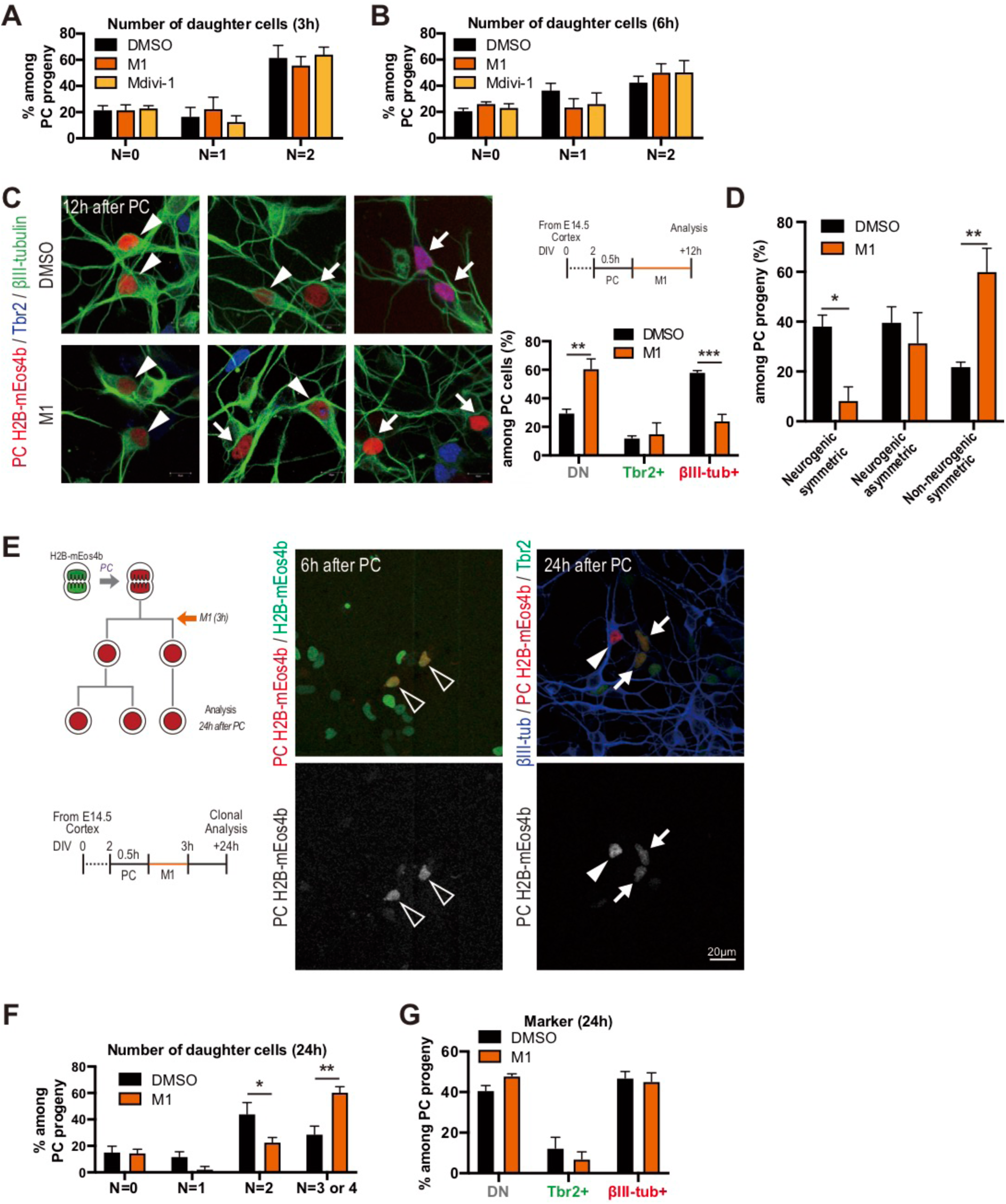
Manipulation of mitochondria dynamics impacts on cortical neurogenesis. (A,B) Quantification of number of daughter cells (A: 3 hours post-PC, B: 6 hours post-PC) from at least four biological replicate experiments. Data are shown as mean ± SEM. (C) Timeline of M1 treatment in 12 hours post-PC experiment. Representative images of PC daughter cells. Arrow head: Neuron (βIII-tub+), Arrow: Progenitor (Tbr2+ or DN). Quantification of each cell fate marker+ cells among PC cells (12 hours post-PC) from three biological replicate experiments. Data are shown as mean ± SEM. **P < 0.01, ***P < 0.001; Bonferroni’s multiple comparisons test. (D) Quantification of each division type among PC cells (12 hours post-PC) from three biological replicate experiments. Data are shown as mean ± SEM. *P < 0.05, **P < 0.01; Bonferroni’s multiple comparisons test. (E) Timeline and representative images of clonal analysis. White open arrow head: PC cells 6 hours post-PC. Arrow head: Neuron (βIII-tub+), Arrow: Progenitor (Tbr2+) 24 hours post-label. (F) Quantification of clone size (24 hours post-PC) from three biological replicate experiments. Data are shown as mean ± SEM. *P < 0.05, **P < 0.01; Bonferroni’s multiple comparisons test. (G) Quantification of cell fate marker-positive cells among PC cells (24 hours post-PC) from three biological replicate experiments. Data are shown as mean ± SEM.

**Fig. S3.**
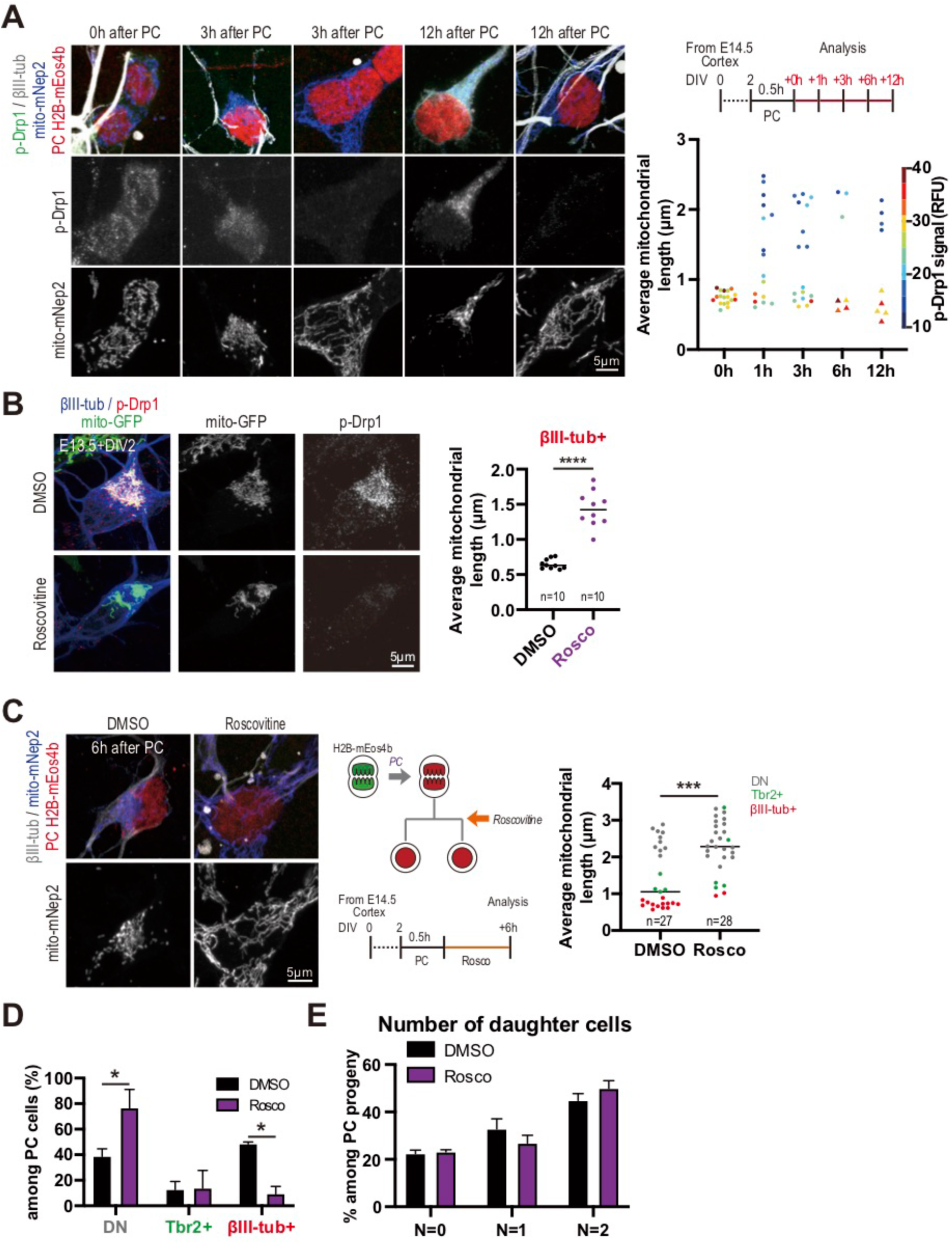
Cdk-dependent mitochondria dynamics of mouse post-mitotic cortical cells. (A) Representative images and timeline of PC experiment to determine mitochondrial morphology (mito-mNep2) together with phospho-Drp1 (p-Drp1) levels. Quantified mitochondrial length and phospho-Drp1 signal intensity (RFU: relative fluorescence unit) from two biological replicate experiments. Each data point represents an individual cell average mitochondrial size and phospho-Drp1 signal. (B) Representative images of mitochondrial morphology and p-Drp1 signal in newborn neurons (E13.5 cortex cultured for 2 DIV). Quantified mitochondrial length of βIII-tub+ from two biological replicate experiments. Each data point represents an individual cell average mitochondrial size. ****P < 0.0001; unpaired t test. (C) Representative images of Roscovitine-treated PC cell and quantified mitochondrial length from three biological replicate experiments. ***P < 0.001; Mann-Whitney test. (D) Quantification of cell fate marker-positive cells among PC cells. Data are shown as mean ± SEM. *P < 0.05; Bonferroni’s multiple comparisons test. (E) Quantification of number of daughter cells from three biological replicate experiments. Data are shown as mean ± SEM.

**Fig. S4.**
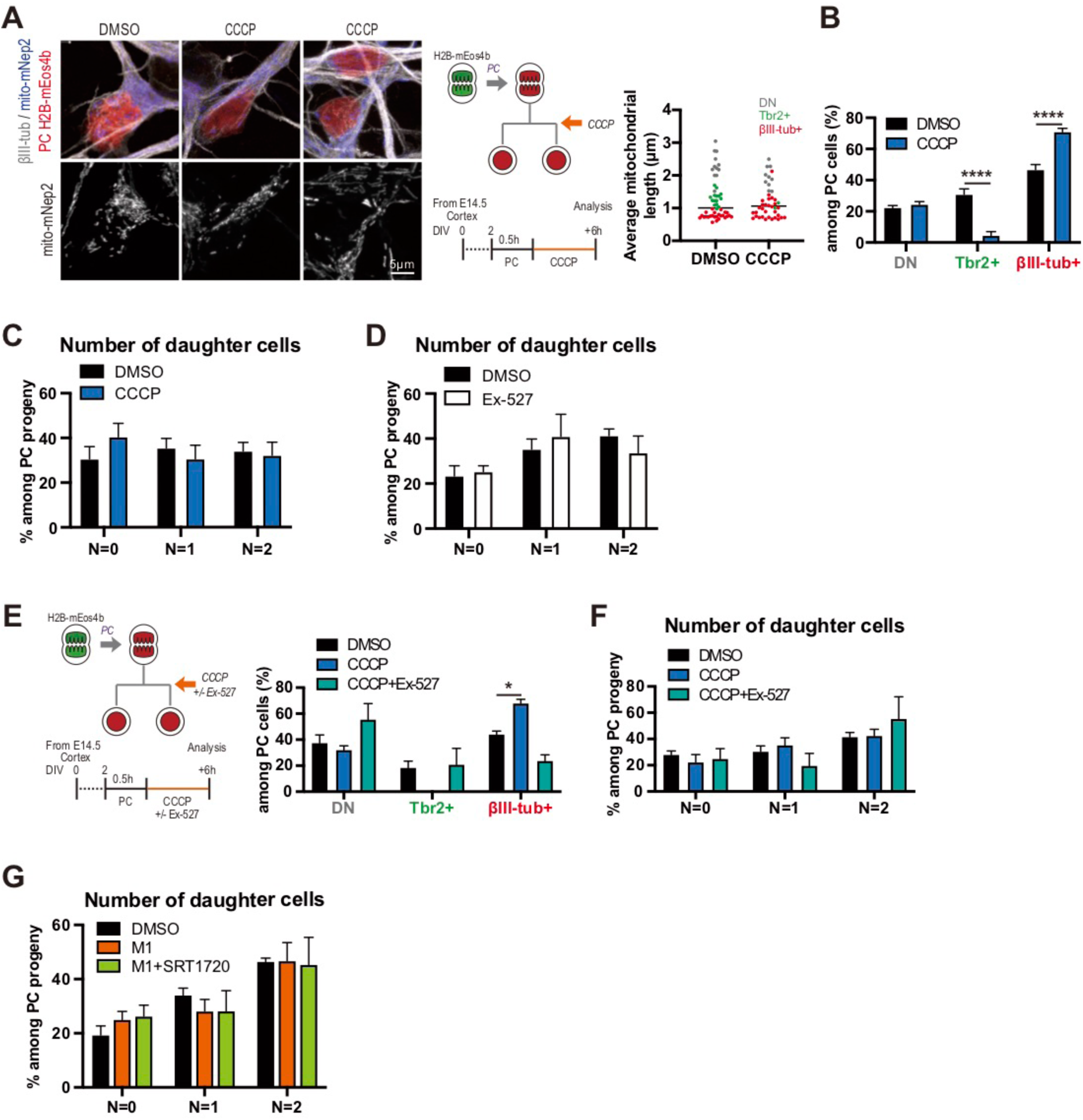
Downstream mechanism of mitochondrial dependent cell fate determination. (A) Timeline, representative images, and quantified mitochondrial length following CCCP postmitotic treatment. Each data point represents an individual cell average mitochondrial size from three biological replicate experiments. Mann-Whitney test. (B) Quantification of each cell fate marker+ cells among PC cells (6 hours post-label) from three biological replicate experiments following CCCP postmitotic treatment. Data are shown as mean ± SEM. ****P < 0.0001; Bonferroni’s multiple comparisons test. (C) Quantification of number of daughter cells under CCCP treatment from three biological replicate experiments. Data are shown as mean ± SEM. (D) Quantification of number of daughter cells under Sirt1 inhibitor Ex-527 treatment from three biological replicate experiments. Data are shown as mean ± SEM. (E) Timeline and quantification of cell fate marker-positive cells among PC cells following CCCP and Ex-527 treatment (6 hours post-PC) from four biological replicate experiments. Data are shown as mean ± SEM, *P < 0.05; Dunnett’s multiple comparisons test. (F) Quantification of number of daughter cells under CCCP and Ex-527 treatments from four biological replicate experiments. Data are shown as mean ± SEM. (G) Quantification of number of daughter cells under M1 and SRT1720 treatments from four biological replicate experiments. Data are shown as mean ± SEM.

**Fig. S5.**
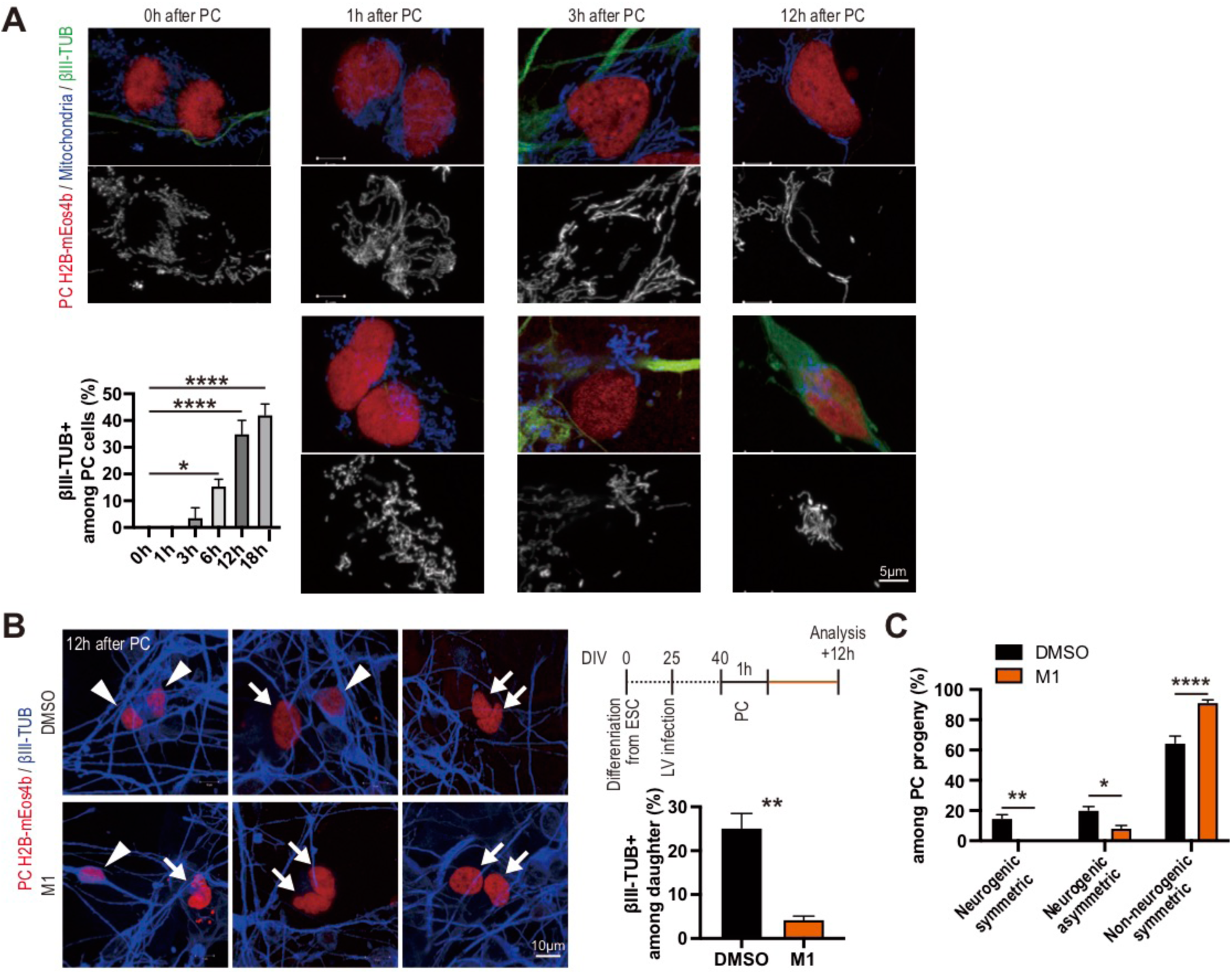
Mitochondrial dynamics and their impacts on cell fate determination in human cortical cells. (A) Representative images of PC cell mitochondrial morphology and cell fate marker in human cortical cells (cf. Fig.4E). Quantification of βIII-TUB+ cells among PC cells from three biological replicate experiments. Data are shown as mean ± SEM. *P < 0.05, ****P < 0.0001; Dunnett’s multiple comparisons test. (B) Timeline and representative images of PC human cortical cells (12 hours post-PC) of M1 treatment in PC strategy. Arrow head: Neuron (βIII-TUB+), Arrow: Progenitor (βIII-TUB–). Quantification of βIII-TUB+ cells among PC cells from three biological replicate experiments. Data are shown as mean ± SEM. **P < 0.01; Unpaired t test. (C) Quantification of cell division type among PC cells (12 hours post-PC) from three biological replicate experiments. Data are shown as mean ± SEM. *P < 0.05, **P < 0.01, ****P < 0.0001; Bonferroni’s multiple comparisons test.

## References

1. C. Lorenz, A. Prigione, Mitochondrial metabolism in early neural fate and its relevance for neuronal disease modeling. Curr. Opin. Cell Biol. (2017), doi:10.1016/j.ceb.2017.12.004.

2. M. Götz, W. B. Huttner, The cell biology of neurogenesis. Nat Rev Mol Cell Biol. 6, 777–788 (2005).

3. D. L. Moore, G. A. Pilz, M. J. Araúzo-Bravo, Y. Barral, S. Jessberger, A mechanism for the segregation of age in mammalian neural stem cells. Science (80-.). (2015), doi:10.1126/science.aac9868.

4. K. Mitra, R. Rikhy, M. Lilly, J. Lippincott-Schwartz, DRP1-dependent mitochondrial fission initiates follicle cell differentiation during Drosophila oogenesis. J. Cell Biol. (2012), doi:10.1083/jcb.201110058.

5. M. D. D. Buck, D. O’Sullivan, R. I. I. Klein Geltink, J. D. D. Curtis, C. H. Chang, D. E. E. Sanin, J. Qiu, O. Kretz, D. Braas, G. J. J. W. van der Windt, Q. Chen, S. C. C. Huang, C. M. M. O’Neill, B. T. T. Edelson, E. J. J. Pearce, H. Sesaki, T. B. B. Huber, A. S. S. Rambold, E. L. L. Pearce, Mitochondrial Dynamics Controls T Cell Fate through Metabolic Programming. Cell. 166, 63–76 (2016).

6. M. Khacho, A. Clark, D. S. Svoboda, J. Azzi, J. G. MacLaurin, C. Meghaizel, H. Sesaki, D. C. Lagace, M. Germain, M. E. Harper, D. S. Park, R. S. Slack, Mitochondrial Dynamics Impacts Stem Cell Identity and Fate Decisions by Regulating a Nuclear Transcriptional Program. Cell Stem Cell. 19, 232–247 (2016).

7. A. Kasahara, S. Cipolat, Y. Chen, G. W. Dorn, L. Scorrano, Mitochondrial fusion directs cardiomyocyte differentiation via calcineurin and notch signaling. Science (80-.). (2013), doi:10.1126/science.1241359.

8. A. Bahat, A. Gross, Mitochondrial plasticity in cell fate regulation. J. Biol. Chem. 294, 13852–13863 (2019).

9. L. Pernas, L. Scorrano, Mito-Morphosis: Mitochondrial Fusion, Fission, and Cristae Remodeling as Key Mediators of Cellular Function. Annu. Rev. Physiol. 78, 505–531 (2016).

10. P. Katajisto, J. Döhla, C. L. Chaffer, N. Pentinmikko, N. Marjanovic, S. Iqbal, R. Zoncu, W. Chen, R. A. Weinberg, D. M. Sabatini, Asymmetric apportioning of aged mitochondria between daughter cells is required for stemness. Science (80-.). 348, 340–343 (2015).

11. K. Ito, R. Turcotte, J. Cui, S. E. Zimmerman, S. Pinho, T. Mizoguchi, F. Arai, J. M. Runnels, C. Alt, J. Teruya-Feldstein, J. C. Mar, R. Singh, T. Suda, C. P. Lin, P. S. Frenette, K. Ito, Self-renewal of a purified Tie2+ hematopoietic stem cell population relies on mitochondrial clearance. Science (80-.). 354, 1156–1160 (2016).

12. N. Bertrand, D. S. Castro, F. Guillemot, Proneural genes and the specification of neural cell types. Nat Rev Neurosci. 3, 517–530 (2002).

13. Y. Hirabayashi, Y. Gotoh, Epigenetic control of neural precursor cell fate during development. Nat Rev Neurosci. 11, 377–388 (2010).

14. B. Yao, K. M. Christian, C. He, P. Jin, G. L. Ming, H. Song, Epigenetic mechanisms in neurogenesis. Nat. Rev. Neurosci. (2016), doi:10.1038/nrn.2016.70.

15. K. Mitra, C. Wunder, B. Roysam, G. Lin, J. Lippincott-Schwartz, A hyperfused mitochondrial state achieved at G1-S regulates cyclin E buildup and entry into S phase. Proc. Natl. Acad. Sci. U. S. A. 106, 11960–11965 (2009).

16. T. L. Lewis, S. K. Kwon, A. Lee, R. Shaw, F. Polleux, MFF-dependent mitochondrial fission regulates presynaptic release and axon branching by limiting axonal mitochondria size. Nat. Commun. (2018), doi:10.1038/s41467-018-07416-2.

17. P. Mishra, D. C. Chan, Mitochondrial dynamics and inheritance during cell division, development and disease. Nat. Rev. Mol. Cell Biol. (2014), doi:10.1038/nrm3877.

18. D. Wang, J. Wang, G. M. C. Bonamy, S. Meeusen, R. G. Brusch, C. Turk, P. Yang, P. G. Schultz, A small molecule promotes mitochondrial fusion in mammalian cells. Angew. Chemie - Int. Ed. 51, 9302–9305 (2012).

19. A. Cassidy-Stone, J. E. Chipuk, E. Ingerman, C. Song, C. Yoo, T. Kuwana, M. J. Kurth, J. T. Shaw, J. E. Hinshaw, D. R. Green, J. Nunnari, Chemical Inhibition of the Mitochondrial Division Dynamin Reveals Its Role in Bax/Bak-Dependent Mitochondrial Outer Membrane Permeabilization. Dev. Cell. 14, 193–204 (2008).

20. N. Taguchi, N. Ishihara, A. Jofuku, T. Oka, K. Mihara, Mitotic phosphorylation of dynamin-related GTPase Drp1 participates in mitochondrial fission. J. Biol. Chem. 282, 11521–11529 (2007).

21. L. Telley, S. Govindan, J. Prados, I. Stevant, S. Nef, E. Dermitzakis, A. Dayer, D. Jabaudon, Sequential transcriptional waves direct the differentiation of newborn neurons in the mouse neocortex. Science (80-.). 351, 1443–1446 (2016).

22. M. Liesa, O. S. Shirihai, Mitochondrial dynamics in the regulation of nutrient utilization and energy expenditure. Cell Metab. 17, 491–506 (2013).

23. N. M. C. Connolly, P. Theurey, V. Adam-Vizi, N. G. Bazan, P. Bernardi, J. P. Bolaños, C. Culmsee, V. L. Dawson, M. Deshmukh, M. R. Duchen, H. Düssmann, G. Fiskum, M. F. Galindo, G. E. Hardingham, J. M. Hardwick, M. B. Jekabsons, E. A. Jonas, J. Jordán, S. A. Lipton, G. Manfredi, M. P. Mattson, B. A. McLaughlin, A. Methner, A. N. Murphy, M. P. Murphy, D. G. Nicholls, B. M. Polster, T. Pozzan, R. Rizzuto, J. Satrústegui, R. S. Slack, R. A. Swanson, R. H. Swerdlow, Y. Will, Z. Ying, A. Joselin, A. Gioran, C. Moreira Pinho, O. Watters, M. Salvucci, I. Llorente-Folch, D. S. Park, D. Bano, M. Ankarcrona, P. Pizzo, J. H. M. Prehn, Guidelines on experimental methods to assess mitochondrial dysfunction in cellular models of neurodegenerative diseases (Springer US, 2018; http://dx.doi.org/10.1038/s41418-017-0020-4), vol. 25.

24. C. Cantó, K. J. Menzies, J. Auwerx, NAD+ Metabolism and the Control of Energy Homeostasis: A Balancing Act between Mitochondria and the Nucleus. Cell Metab. 22, 31–53 (2015).

25. X. R. Bao, S. E. Ong, O. Goldberger, J. Peng, R. Sharma, D. A. Thompson, S. B. Vafai, A. G. Cox, E. Marutani, F. Ichinose, W. Goessling, A. Regev, S. A. Carr, C. B. Clish, V. K. Mootha, Mitochondrial dysfunction remodels one-carbon metabolism in human cells. Elife. 5, 1–24 (2016).

26. L. Tiberi, J. Van Den Ameele, J. Dimidschstein, J. Piccirilli, D. Gall, A. Herpoel, A. Bilheu, J. Bonnefont, M. Iacovino, M. Kyba, T. Bouschet, P. Vanderhaeghen, BCL6 controls neurogenesis through Sirt1-dependent epigenetic repression of selective Notch targets. Nat Neurosci. 15, 1627–1635 (2012).

27. S. Hisahara, S. Chiba, H. Matsumoto, M. Tanno, H. Yagi, S. Shimohama, M. Sato, Y. Horio, Histone deacetylase SIRT1 modulates neuronal differentiation by its nuclear translocation. Proc Natl Acad Sci U S A. 105, 15599–15604 (2008).

28. J. Bonnefont, L. Tiberi, J. van den Ameele, D. Potier, Z. B. Gaber, X. Lin, A. Bilheu, A. Herpoel, F. D. Velez Bravo, F. Guillemot, S. Aerts, P. Vanderhaeghen, Cortical Neurogenesis Requires Bcl6-Mediated Transcriptional Repression of Multiple Self-Renewal-Promoting Extrinsic Pathways. Neuron. 103 (2019), doi:10.1016/j.neuron.2019.06.027.

29. J. H. Lui, D. V. Hansen, A. R. Kriegstein, Development and evolution of the human neocortex. Cell (2011), doi:10.1016/j.cell.2011.06.030.

30. A. M. M. Sousa, K. A. Meyer, G. Santpere, F. O. Gulden, N. Sestan, Evolution of the Human Nervous System Function, Structure, and Development. Cell (2017), doi:10.1016/j.cell.2017.06.036.

31. I. K. Suzuki, D. Gacquer, R. Van Heurck, D. Kumar, M. Wojno, A. Bilheu, A. Herpoel, N. Lambert, J. Cheron, F. Polleux, V. Detours, P. Vanderhaeghen, Human-Specific NOTCH2NL Genes Expand Cortical Neurogenesis through Delta/Notch Regulation. Cell. 173, 1370–1384 e16 (2018).

32. I. Espuny-camacho, K. A. Michelsen, D. Gall, D. Linaro, A. Hasche, N. Lambert, N. Gaspard, S. Pe, C. Bali, D. Orduz, S. N. Schiffmann, M. Giugliano, A. Gaillard, P. Vanderhaeghen, J. Bonnefont, C. Bali, D. Orduz, A. Bilheu, A. Herpoel, N. Lambert, N. Gaspard, S. Péron, S. N. Schiffmann, M. Giugliano, A. Gaillard, P. Vanderhaeghen, Pyramidal Neurons Derived from Human Pluripotent Stem Cells Integrate Efficiently into Mouse Brain Circuits In Vivo. Neuron. 77, 440–456 (2013).

33. C. Dehay, H. Kennedy, Cell-cycle control and cortical development. Nat Rev Neurosci. 8, 438–450 (2007).

34. P. Salomoni, F. Calegari, Cell cycle control of mammalian neural stem cells: Putting a speed limit on G1. Trends Cell Biol. 20, 233–243 (2010).

35. T. Takahashi, R. S. Nowakowski, V. S. Caviness, The cell cycle of the pseudostratified ventricular epithelium of the embryonic murine cerebral wall. J. Neurosci. (1995), doi:10.1523/jneurosci.15-09-06046.1995.

36. T. Edlund, T. M. Jessell, Progression from extrinsic to intrinsic signaling in cell fate specification: A view from the nervous system. Cell. 96, 211–224 (1999).

37. S. Dalton, Linking the Cell Cycle to Cell Fate Decisions. Trends Cell Biol. 25, 592–600 (2015).

38. L. M. Julian, R. L. Carpenedo, J. L. M. Rothberg, W. L. Stanford, Formula G1: Cell cycle in the driver’s seat of stem cell fate determination. BioEssays. 38, 325–332 (2016).

39. H. Zhang, D. J. Emerson, T. G. Gilgenast, K. R. Titus, Y. Lan, P. Huang, D. Zhang, H. Wang, C. A. Keller, B. Giardine, R. C. Hardison, J. E. Phillips-Cremins, G. A. Blobel, Chromatin structure dynamics during the mitosis-to-G1 phase transition. Nature. 576, 1–5 (2019).

40. R. Beckervordersandforth, B. Ebert, I. Schäffner, J. Moss, C. Fiebig, J. Shin, D. L. Moore, L. Ghosh, M. F. Trinchero, C. Stockburger, K. Friedland, K. Steib, J. von Wittgenstein, S. Keiner, C. Redecker, S. M. Hölter, W. Xiang, W. Wurst, R. Jagasia, A. F. Schinder, G. li Ming, N. Toni, S. Jessberger, H. Song, D. C. Lie, Role of Mitochondrial Metabolism in the Control of Early Lineage Progression and Aging Phenotypes in Adult Hippocampal Neurogenesis. Neuron. 93, 560–573.e6 (2017).

41. N. Shyh-Chang, G. Q. Daley, L. C. Cantley, Stem cell metabolism in tissue development and aging. Dev. 140, 2535–2547 (2013).

42. C. C. F. Homem, M. Repic, J. A. Knoblich, Proliferation control in neural stem and progenitor cells. Nat. Rev. Neurosci. 16, 647–659 (2015).

43. M. Knobloch, S. Jessberger, Metabolism and neurogenesis. Curr. Opin. Neurobiol. 42, 45–52 (2017).

44. X. Zheng, L. Boyer, M. Jin, J. Mertens, Y. Kim, L. Ma, L. Ma, M. Hamm, F. H. Gage, T. Hunter, Metabolic reprogramming during neuronal differentiation from aerobic glycolysis to neuronal oxidative phosphorylation. Elife. 5, 1–25 (2016).

45. M. Agathocleous, N. K. Love, O. Randlett, J. J. Harris, J. Liu, A. J. Murray, W. A. Harris, Metabolic differentiation in the embryonic retina. Nat. Cell Biol. 14, 859–864 (2012).

46. H. Shimojo, T. Ohtsuka, R. Kageyama, Oscillations in notch signaling regulate maintenance of neural progenitors. Neuron. 58, 52–64 (2008).

47. J. G. Ryall, S. Dell’Orso, A. Derfoul, A. Juan, H. Zare, X. Feng, D. Clermont, M. Koulnis, G. Gutierrez-Cruz, M. Fulco, V. Sartorelli, The NAD+-dependent sirt1 deacetylase translates a metabolic switch into regulatory epigenetics in skeletal muscle stem cells. Cell Stem Cell (2015), doi:10.1016/j.stem.2014.12.004.

48. S. Gascón, G. Masserdotti, G. L. Russo, M. Götz, Direct Neuronal Reprogramming: Achievements, Hurdles, and New Roads to Success. Cell Stem Cell. 21 (2017), pp. 18–34.

